# Honestly exaggerated: howler monkey roars are reliable signals of body size and behaviourally relevant to listeners

**DOI:** 10.1101/2024.12.06.626400

**Authors:** Jacob C. Dunn, Eloise Pederson, Holly Farmer, Philippa Dobbs, W. Tecumseh Fitch, David Reby, Benjamin Charlton

**Author notes:** Correspondence to: Dr Jacob C. Dunn, Behavioural Ecology Research Group, Anglia Ruskin University, East Road, Cambridge, CB1 1PT.

## Abstract

Acoustic signals are key components of animal social behaviour, potentially conveying fitness relevant information about signallers. Howler monkeys produce extremely loud, low frequency roars, which exaggerate the acoustic impression of body size relative to other species. However, it remains unclear whether howler monkey roars contain reliable information about body size within species, and whether conspecific listeners use this information and adjust their responses accordingly. Here, we investigate whether the roars of black and gold howler monkeys (*Alouatta caraya*) function as an honest signal of body size by examining the relationship between formant spacing and body mass in 11 adult males. We found a strong negative correlation, indicating that larger males produce roars with lower formant spacing. To test the behavioural relevance of variation in formant spacing, we then conducted playback experiments with 23 conspecific listeners, simulating the roars of unknown males of small, average, and large body size. Listeners showed significantly different responses to calls of different body sizes, spending longer orientated towards the playback speaker and being more likely to approach calls simulating larger males. There was no significant impact of simulated body size on the likelihood of listeners vocalising in response, although males spent significantly more time vocalising in response to playbacks than females. Overall, these findings suggest that formant spacing in howler monkey roars serves as an honest indicator of body size and plays a critical role in mediating social interactions. Our study highlights the adaptive significance of acoustic cues to body size, which can provide receivers with accurate information that can be used to assess rivals or choose mates.

## Introduction

Large animals typically produce lower frequency calls than smaller animals, for the simple reason that they generally have larger vocal tracts and/or vocal sources. The negative correlation between call frequency and body size (or ‘negative acoustic allometry’), has been recognised for decades, and provides the underpinnings of the idea of ‘honest’ advertising (Bowling et al., 2020, 2017; Clutton-Brock & Albon, 1979; Davies & Halliday, 1978; Fletcher, 2004; Garcia et al., 2017; Martin et al., 2017; Morton, 1977; Reby & McComb, 2003). Honest calls are postulated to provide reliable information to receivers about important attributes of the signaller, e.g., sex, age, reproductive status, and motivation (Fitch & Hauser, 1995, 1998; Gouzoules & Gouzoules, 2002; Suthers et al., 2016; Taylor & Reby, 2010).

Research has frequently focussed on the link between body size and mean and/or lowest fundamental frequency (*f*_o_). The lowest fundamental frequency (*f*_o_min) is determined by the length and mass of the vocal folds – longer vocal folds produce a lower *f*_o_min (Riede & Brown, 2013; Titze et al., 2016; Titze, 1994). Thus, across species, a negative relationship between *f*_o_min and body size has been described (Bowling et al., 2017; Fletcher, 2004; Garcia et al., 2017; Morton, 1977; Tembrock, 1996), and the size of the vocal folds (and thus of *f*_o_) could, in principle, provide information about body size to receivers, which may regulate their behaviour accordingly (e.g., towards potential mates or competitors). This also appears to be the case across individuals within certain taxa, including some frogs and toads (Davies & Halliday, 1978; Martin, 1972; Ryan, 1988). However, in other vertebrate taxa, there is no reliable relationship between *f*_o_ and body size within a species. For example, there is no correlation between *f*_o_ and body size in adult humans, within either sex (Cohen et al., 1980; Lass, 1978; Pisanski et al., 2014; van Dommelen, 1993; van Dommelen & Moxness, 1995). Similarly, in many species of birds (Ryan & Brenowitz, 1985), amphibians (Asquith & Altig, 1990), and mammals (Bowling et al., 2017; Charlton & Reby, 2016; Garcia et al., 2017; Masataka, 1994; McComb, 1991; Rendall et al., 2005), there appears to be no relationship between body size and *f*_o_.

The lack of a clear relationship between body size and *f*_o_ is linked to the underlying anatomy of the larynx and vocal folds. The larynx is a cartilaginous organ, which is not tightly constrained in size by its neighbouring bony structures (Fitch 1997; Harrison, 1995; Negus, 1962). Therefore, the larynx and vocal folds can grow somewhat independently of the head and the rest of the body. In human males, for example, the vocal folds lengthen at puberty, lowering *f*_o_, and vocal fold length is thought to link to testosterone levels (Fouquet et al., 2016; Kahane, 1978; Kreiman & Sidtis, 2011; Markova et al., 2016; Titze, 1994). Given this decoupling between body size and the size of the larynx, there is no *a priori* reason why we should expect a strong relationship between *f*_o_ and body size within species.

Recent research has instead focussed on formant frequencies, the resonances of the vocal tract which produce spectral peaks when the vocal tract filters the sound produced within the larynx (Fant, 1960; Fitch, 1997; Fitch & Hauser, 1995; Taylor & Reby, 2010; Titze, 1994). The formant frequencies (F_1_ – F_n_) are determined by the length and shape of the vocal tract (the air-filled chambers of the throat, nose, and mouth). Larger animals, with longer vocal tracts, are expected to produce lower formant frequencies than smaller animals. This prediction has found to be more consistently supported than that between body size and *f*_o_. In particular, formant spacing (the average spacing between formants; (Fitch, 1997; Reby & McComb, 2003; Taylor & Reby, 2010; Titze, 1994) has been found to be strongly linked to body size across a range of taxa (Charlton et al., 2009; Charlton & Reby, 2016; Fitch, 1997; Pisanski et al., 2014; Reber et al., 2017; Reby & McComb, 2003; Riede & Fitch, 1999; Vannoni & McElligott, 2008). This is likely because vocal tract length is much more constrained by body size than vocal fold length (Fitch, 1997).

Some species have evolved anatomical adaptations to exaggerate the acoustic advertisement of body size (Fitch & Reby, 2001; Taylor et al., 2016). In some cases, this seems to be achieved through adaptations in the larynx and vocal folds, impacting on *f*_o_. For example, in humans and red deer, the vocal folds are sexually dimorphic, lengthening in males at puberty. Males in some species have extremely large larynges for their body size, such as hammerhead bats (Schneider et al., 1967). An extreme case of low *f*_o_ sound production is observed in the koala, which has evolved a second pair of ‘velar vocal folds’, which are substantially longer than the laryngeal vocal folds, and produce very low frequency sound (Charlton et al., 2013).

Nevertheless, while *f*_o_ may provide an estimate of body size to listeners *across* species (or even across different age and/or sex classes within species), it is usually not a reliable signal of body size *within* a species or age classes, for the reasons discussed above. An alternative means of increasing the acoustic impression of body size would be to increase vocal tract size/length, determining formant dispersion. This is likely to be a source of much more reliable (or ‘honest’) information about body size. For example, some mammal species possess a ‘descended larynx’ which lengthens the vocal tract (reviewed in (Taylor et al., 2016). Similarly, some cervids are able to retract the larynx during vocalisation, allowing them to lengthen the vocal tract dynamically, and thus lower formants (Fitch & Reby, 2001). Other species have evolved extensions to the vocal tract which decrease formant dispersion. For example, saiga antelopes (Frey et al., 2007), elephants (McComb et al., 2003) and elephant seals (Sanvito et al., 2007) all use an extended proboscis during vocalisation. Similarly, many primate species have evolved air sacs which act as additional resonators during vocalisation (de Boer, 2009; Fitch & Hauser, 1995; Harris et al., 2006).

In a few species, such adaptations are taken to extremes. For example, howler monkeys (*Alouatta* spp.) have evolved a hypertrophied larynx, out of all proportion to body size, in which the larynx and hyoid are bigger than the entire cranium in some species (Dunn et al., 2015; Schön, 1971). The result is that an 8kg male howler monkey has the vocal apparatus equivalent in size to that of a 500kg Siberian tiger, and produces roars with extremely low *f*_o_ and formant dispersion (Dunn et al., 2015). Indeed, howler monkey calls appear to predict a vocal tract length of up to 45 cm, longer than the total body length of only about 40–50 cm in this genus (Dunn et al., 2015).

Such calls have likely evolved in howler monkeys to exaggerate the acoustic impression of the calling individual’s body size. Although these acoustic cues appear to be exaggerated relative to other species, it remains unclear whether such calls contain reliable information about body size for conspecifics across howler species (but see (De los Santos Mendoza & Van Belle, 2024). Even an exaggerated trait is likely to ultimately be limited by some anatomical constraint, and if these are linked to body size, then this would generate a signal that would function honestly within the species, although exaggerated relative to other species. For example, the retractable larynx of red deer is limited in its descent by the sternum, and in extreme roars (termed “great roars”), is unable to descend any further. Thus, although red deer calls acoustically exaggerate body size, bigger individuals still sound bigger than smaller individuals, as they have a longer vocal tract (Reby & McComb, 2003).

Here, we evaluate the extreme vocal adaptations found in howler monkeys to determine whether their loud, low frequency calls contain honest information about body size. We then use playback experiments to evaluate whether howler monkeys perceive variation in conspecific formant dispersion. We test two predictions: 1) that formant dispersion is negatively correlated with body size; and 2) that conspecific howler monkeys respond more strongly to acoustic stimuli which simulate larger individuals than smaller individuals.

## Methods

### Study species

Black-and-gold howler monkeys (*Alouatta caraya*) are naturally found in the forests of Argentina, Bolivia, Brazil, Paraguay and Uruguay (Cunha et al., 2015). They live in uni- or multi-male groups, with an average group size of 10 individuals (Dunn et al., 2015). They roar loudly at dawn and dusk in order to regulate space by means of regular advertisement of territory occupancy (Cunha & Byrne, 2006). Their sexually dimorphic vocal apparatus suggests that sexual selection is likely to drive the evolution of vocal communication, with males producing significantly lower frequency calls than females (Dunn et al., 2015). Howler monkeys are generally very sensitive to captive environments, and most species are unable to tolerate captivity (Pastor-Nieto, 2015). However, black-and-gold howler monkeys are found in numerous zoos across the world and provide ideal study subjects to test the hypotheses advanced here.

### Recordings and body size data

Recordings of adult male roars were made from 11 captive male howler monkeys. This included 8 individuals housed at 5 different zoos across the UK and 3 individuals housed in 3 different zoos in the USA (Table S1). The roars in the UK were recorded with a solid-state recorder and a directional microphone. For the 3 individuals in the USA, recordings were taken with a video camera and internal microphone. Recordings were made at a sampling frequency of 44.1 kHz and 24-bit accuracy.

The most recent data on body size for the date of the recording were provided for the 11 males by the veterinary staff at the zoos. We used body mass (kg) as our proxy for body size, as this is the measurement that was consistently recorded across all zoos. Data from Twycross Zoo show that across 43 adult individuals, body mass (kg) is strongly correlated with crown-rump length (cm) (y = 11.39ln(x) +18.26; R² = 0.79). Therefore, body mass appears to be a reliable measure of overall body size, and none of the individuals were considered overweight.

### Acoustic analysis

Acoustic analyses were conducted using the Praat 6.0.31 DSP package (www.praat.org) (Boersma & Weenink, 2001). Six prominent frequency components exist below 3,200 Hz in the spectral acoustic structure of roars, which are likely to represent formants (Dunn et al., 2015). Accordingly, we measured the frequency values (Hz) of the first six spectral prominences (hereafter formants; F1-F6) using Linear Predictive Coding (‘To Formants (Burg)’ command in Praat). We did not measure formant frequencies above 3200 Hz because they were often poorly defined.

The minimum frequency values of each of the first six formant frequencies were extracted from each recording and measured using the following analysis parameters: time step: 0.01 seconds; window analysis: 0.03 seconds; pre-emphasis: 50 Hz, maximum number of formants: 6; maximum formant value: 3200 Hz). To verify that Praat was accurately tracking formants we compared the outputs with visual inspections of each call’s spectrogram and power spectrum (using cepstral smoothing: 200 Hz). Because roars are produced with an open mouth (de Cunha et al., 2015; Kitchen et al., 2015; Schön Ybarra, 2008) we modelled the vocal tract as a tube open at one end (the mouth) and closed at the other (the glottis). A linear tube open at one end and closed at the other acts as a quarter-wave resonator, which should produce a first formant frequency (F1) at c/4L (in which L = length and c is the speed of sound: 350 metres per second in the moist air of the vocal tract), and subsequent formants at F1*3, F1*5, F1*7 etc. (Titze, 1994). Using the ‘open-one-end’ vocal tract model we then estimated the formant spacing (ΔF) during each roar using a linear regression model (Reby & McComb, 2003).

### Playback sequences

We constructed playback sequences from recordings of wild adult male black and gold howler monkeys taken from the Macaulay Library and British Library. Thus, these individuals were previously unknown to the zoo subjects. More than 20 recordings of this species were screened, and four recordings of similar length were chosen, optimising the quality of the recordings (e.g., signal to noise ratio).

We re-synthesised these 4 roars to create large and small variants with ΔF increased and decreased by 5% from the original recordings. To determine the appropriate percentage by which to re-synthesise the playback stimuli, we examined previously reported variation in ΔF in black and gold howler monkeys (Dunn et al., 2015), as well as the variation observed in our own sample of recordings from the 11 captive males. A 5% increase/decrease was used (Fig. 1) to represent the approximately 10% natural variation in adult ΔF in both of these samples.

**Figure 1.**
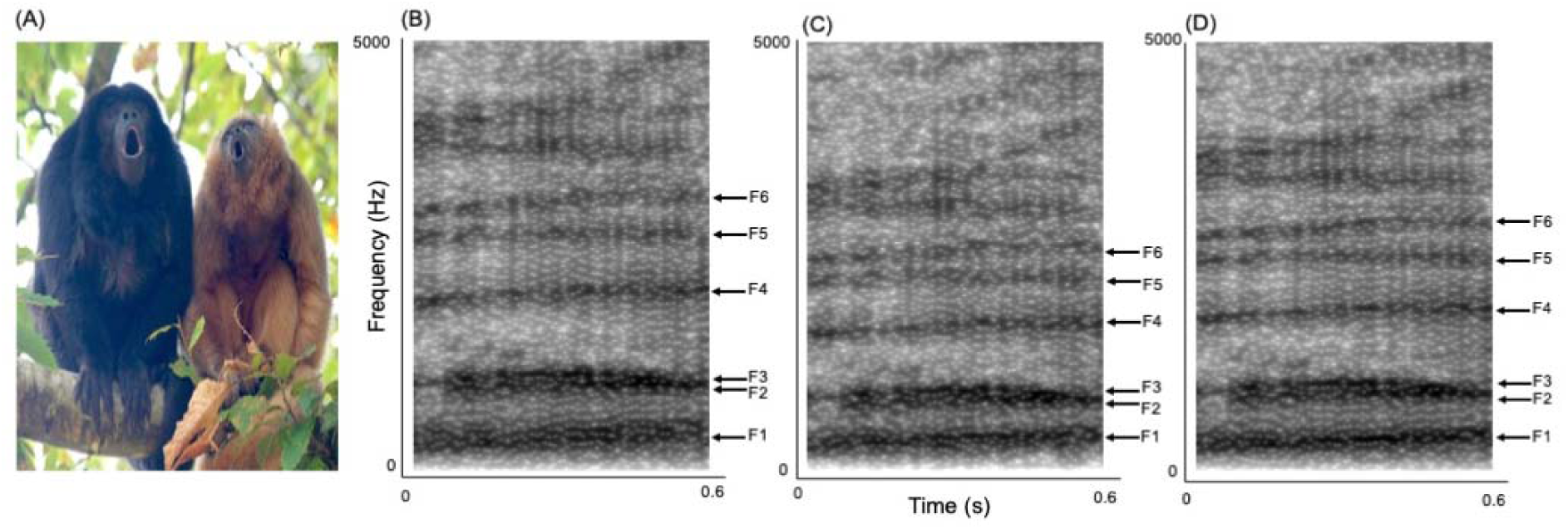
A) Male (black) and female (gold) howler monkeys roaring (© Jacob Dunn); B-D) Formant modification: the spectrograms (settings: FFT method; window length 0.05 s; time step 0.004 s; frequency step 20 Hz; Gaussian window shape; dynamic range 70 dB) show a section of a male roar with (B) small, (C) large, and (D) normal size variants. Note that the formants (labelled F1-F6) have been raised to create the small size variant and lowered to create the large size variant.

Roar modification was completed using a PSOLA-based algorithm that shifts ΔF by a factor (k) while leaving all other acoustic parameters unchanged (for more details see (Charlton, Ellis, et al., 2012; Charlton et al., 2008, 2010)). The recordings were edited to be of comparable duration (70.5 ± 7.7 sec) and the mean relative intensity values for all the playback stimuli were standardized using the ‘scale intensity’ command in Praat.

### Playback experiments

Playback subjects included a total of 23 individuals living in five groups of black-and-gold howler monkeys at Twycross Zoo and Port Lympne. Recordings were broadcast through a MiPro MA707 70W Wireless PA speaker elevated at a height of 3m from the ground and connected to an iPad. Presenting the stimuli at a height of 3m allowed us to limit the effect of ground reflections and was considered more realistic for simulating an arboreal primate to animals living in an enclosure of limited height. The speakers were hidden from view and placed 10m away from each group’s outdoor enclosure, with the position of the first playback being either to the left or right of the enclosure, and the position of the second playback being the opposite side. This reduced the chances of the subjects becoming habituated to a call from a specific position. Roars were broadcast from the speaker at sound pressure levels equivalent to that of naturally roaring males (90 dB mean sound pressure level at 1m from the source (Whitehead, 1989), which were determined using a digital sound level meter.

Each group was subjected to 9 playbacks over the course of the study, with 3 calls being original unmodified calls, 3 being modified to have higher ΔF (small male variant), and 3 being modified to have lower ΔF (large male variant). A maximum of two calls were played to each group per day (one in the morning and one in the afternoon), to reduce any potential stress created by hearing a simulation of a conspecific entering their territory, and to reduce habituation. In addition, the calls selected for each day were chosen so the same group would not be subjected to two versions of the same individual on the same day. The playbacks were initiated when the subjects were inactive, but not asleep, and their attention was directed away from the speaker position. This ensured a standardised context at the onset of each experiment.

Behavioural responses to the playback experiments were captured using a Sony DCR-HC62 Megapixel LCD Handy-cam pointing directly at the enclosure from a central position.

### Behavioural Analysis

Behaviours of all group members were examined for the first 3 minutes after the playback was started. This was to ensure that the behaviours were in immediate response to the playback recordings. Specifically, we measured the time spent oriented towards the speaker, whether the individual moved towards the speaker, and whether the individual vocalised in response to playback. Videotapes were analysed with the observer blind to the condition. Orientation towards the speaker was defined as starting when a subject turned and/or raised their head to face the speaker, and ended when the head moved away from this position. Movement towards the speaker was defined as starting when a subject’s entire body moved in a direction that reduced the distance between their current position and the speaker, and ended when the subject stopped moving, or their entire body moved in a direction that increased the distance between their current position and the speaker. Vocalising included any vocalisation produced by a focal group member during the playback experiment.

### Ethical note

Research and ethical approval were granted for the project by the Twycross Zoo Research Committee, the Aspinall Foundation at Port Lympne, and the University of XXXXX Department of XXXXX. Protocols followed ARRIVE (Sert et al., 2020) and ASAB guidance (ASAB Ethical Committee/ABS Animal Care Committee, 2023). Care was taken to minimise any adverse impacts on the welfare of subjects by minimising the number of exposures to 2 playbacks per group per day (one in the morning and one in the afternoon) and playing back stimuli at natural amplitudes. In *Alouatta caraya,* exposure to conspecific roaring has previous been shown to stimulate natural vocalisations, increase reproductive success, and is generally considered to be beneficially for captive welfare (Farmer, 2011; Farmer et al., 2011).

### Data analysis

Statistical analyses were conducted using R v3.2.3 (R Core Team, 2020). Log_10_ transformations were used to convert our measure of male body weight to scale linearly with vocal tract length (since mass should increase as the cube of length). A linear regression was then used to examine the relationship between ΔF and log_10_ male body weight. We used repeated measures generalized estimating equations (GEE) (“geepack” package in R; (Halekoh et al., 2006) to examine the playback data, with the behavioural responses to the playback presentations as dependant variables, and playback condition and sex as predictor variables. Movement towards and vocalising in response to playback were coded as binomial (yes or no) dependant variables.

## Results

We found a strong negative acoustic allometry between log_10_ male body weight and ΔF (N = 11, F_1, 9_ = 14.1, p = 0.005) (Fig. 2A; Table S1).

**Figure 2.**
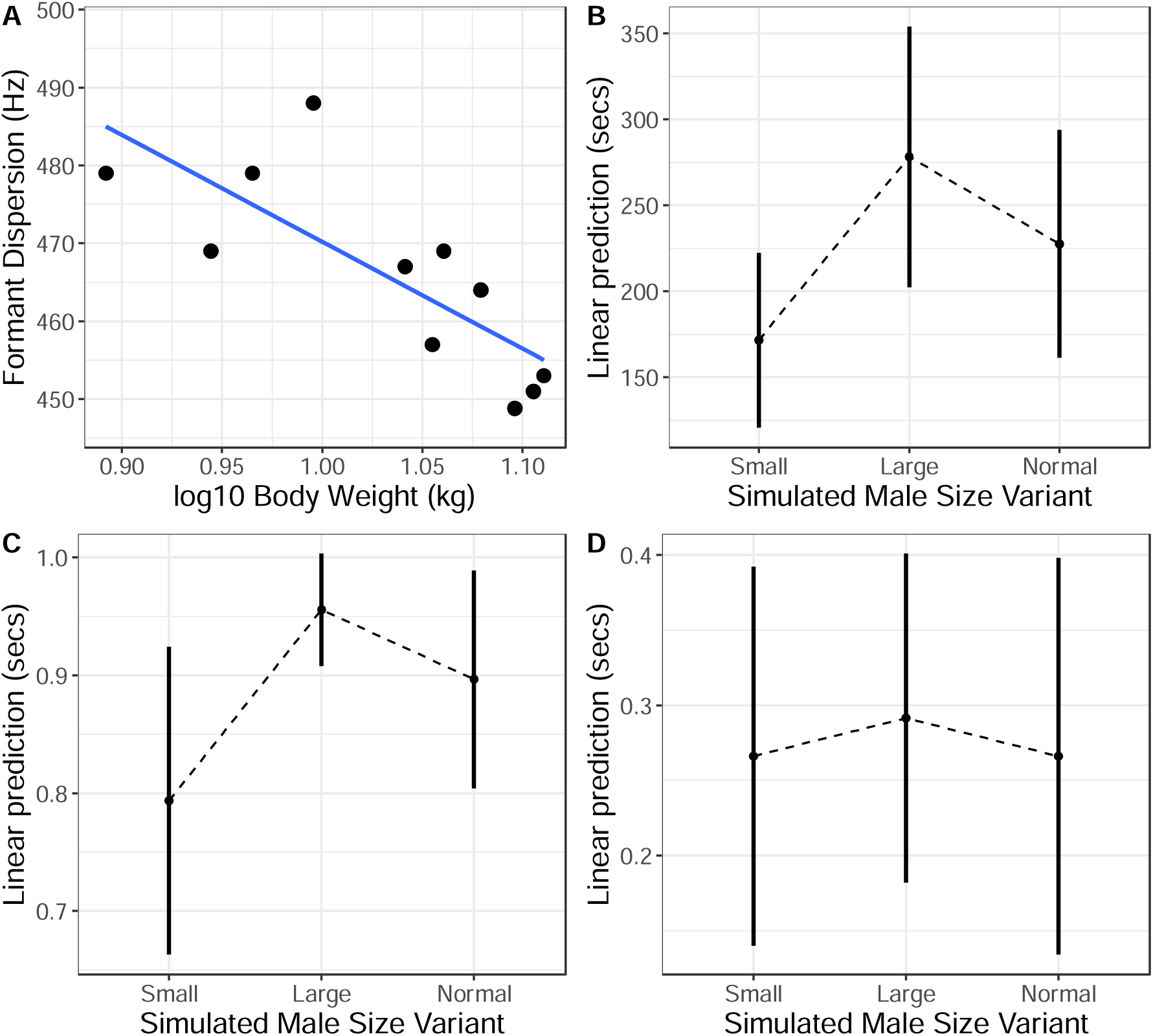
(A) The relationship between formant spacing (Hz) and log_10_ body weight (kg); (B-D). Estimated marginal means ± 95% confidence intervals of behavioural responses to small, large and normal male playback stimuli: (B) time spent orientating towards speaker; (C) likelihood of approaching speaker; (D) likelihood of vocalizing.

Playback condition had a significant effect on the time spent oriented towards (Wald χ^2^_2, 21_ = 54.5, p < 0.001) and likelihood of approaching the speaker (Wald χ^2^_2, 21_ = 9.3, p = 0.010). There was no significant difference between the sexes in time spent orientated towards the speaker (Wald χ^2^_2, 21_ = 0.0, p = 0.880) or likelihood of approach (Wald χ^2^_2, 21_ = 0.0, p = 0.854). Paired comparisons revealed that subjects spent longer oriented towards the large variant than medium (Wald χ^2^_2, 21_ = 7.5, p < 0.006) and small male size variants (Wald χ^2^_2, 21_ = 53.5, p < 0.001) (Fig. 2B). Subjects also spent longer oriented towards speakers broadcasting medium versus small male size variants (Wald χ^2^_2, 21_ = 16.0, p < 0.001) (Fig. 2B). In addition, subjects were more likely to approach speaker positions when the large male condition was presented than the medium (Wald χ^2^_2, 21_ = 4.2, p = 0.042) or small male condition (Wald χ^2^_2, 21_ = 9.21, p = 0.002) (Fig. 3C), and there was an increased tendency for subjects to approach the medium than the small male size condition (Wald χ^2^_2, 21_ = 3.3, p = 0.069) (Fig. 2C). Playback condition was not a significant predictor of the likelihood of vocalizing in response to stimuli (Wald χ^2^_2, 21_ = 0.3, p = 0.882) (Fig 2C). However, an influence of sex on the likelihood of vocalizing was revealed (Wald χ^2^_2, 21_ = 8.4, p = 0.004), with males spending more time vocalising in response to playbacks than females. Raw data from the playback experiments are provided in Table S2.

## Discussion

We found that larger male howler monkeys produced roars with lower formants and formant frequency spacing than smaller males. These findings are in line with our predictions, and accord with research on other mammals, in which formant frequency spacing has been shown to negatively correlate with male body size across a range of mammals (Charlton et al., 2009; Charlton & Reby, 2016; Fitch, 1997; Harris et al., 2006; Reby & McComb, 2003; Riede & Fitch, 1999; Sanvito et al., 2007; Vannoni & McElligott, 2008), and one other species of howler monkey, *Alouatta pigra* (De los Santos Mendoza & Van Belle, 2024). In the current study the mean ΔF of roars from 11 males was 466 Hz (range = 449 to 488 Hz). Assuming that the vocal tract of this species can be modelled as a uniform tube closed at the glottis and opened at the mouth, this value of formant spacing corresponds to a vocal tract length of ∼37 cm, which is much longer than the vocal tract length of 20.6 cm obtained from MRI scans of a male black howler monkey cadaver (Dunn et al., 2015). Formant spacing therefore appears to act as an exaggerated, but honest, signal of body size in black howler monkeys relative to other conspecifics.

Black howler monkeys do not appear to possess anatomical adaptations that would allow them to elongate the supra-laryngeal vocal tract or nasal vocal tract and lower formants, such as a descended larynx (Charlton et al., 2011; Fitch & Reby, 2001; Frey & Gebler, 2003) or nasal proboscis (Frey et al., 2007; Koda et al., 2018; Stoeger et al., 2012). However, the voluminous air sac contained within the greatly enlarged hyoid bone bulla in this species could act as a cavity resonator during vocal production (Kelemen & Sade, 1960; Schön, 1971; Shearer et al., 2015), effectively elongating the vocal tract (de Boer, 2009). Consistent with this notion, howler monkey species with larger hyoids, and presumably a larger hyoid bulla resonator, produce roars with lower formants (Dunn et al., 2015). In addition, because the volume of the hyoid bulla is positively correlated with skull size in black howler monkeys (Dunn et al., 2015) and ultimately constrained by surrounding bone, the formant spacing would remain an honest cue to body size within this species, even if it exaggerates body size relative to other species (similar to findings in red deer and colobus monkeys (Harris et al., 2006; Reby & McComb, 2003). Future studies should develop vocal and nasal tract models using computer tomography scan data (Bowling et al., 2020, 2017; Gamba et al., 2012, 2017; Garcia et al., 2017; Garcia & Dunn, 2019; Ma et al., 2016; Nakamura et al., 2024; Nishimura et al., 2022; Reby et al., 2018) to determine whether the combined area functions from the hyoid bulla and vocal/nasal tract produce the same formant pattern observed in male roars. Close concordance would suggest that selection pressures to broadcast size-related formant information led to the enlargement and pneumatization of this species’ hyoid bone.

Our playback experiments confirmed that male and female black howler monkeys respond differently to large, medium, and small male size variants based on ΔF. This type of spontaneous formant perception, as opposed to that displayed by animals trained to respond to formant variation, provides the strongest indication that formants have functional relevance in a species’ vocal communication system. A number of other nonhuman animals have been shown to spontaneously respond to formant shifts in their own species-typical vocalisations (Charlton, Ellis, et al., 2012; Charlton et al., 2008, 2010; Fitch & Fritz, 2006; Fitch & Kelley, 2000; Reby et al., 2005) and are also capable of perceiving formant shifts in human speech sounds with a high degree of accuracy (Baru, 1975; Burdick & Miller, 1975; Dooling & Brown, 1990; Sinnott, 1989). Playback experiments that test the function of formant variation in species-specific calls, however, are rare in primates, and so far restricted to one genus, *Macaca* (rhesus macaque, *Macaca mulatta*, (Fitch & Fritz 2006, Ghazanfar et al., 2007); Japanese macaque, *Macaca fuscata*, (Furuyama et al., 2016).

In the current study we have shown that male and female black howler monkeys adjust their behavioural responses according to the size-related information broadcast in male roars, paying more attention and approaching more towards speakers broadcasting roars simulating larger male callers. We suggest the most plausible explanation for these findings is that the larger size variant represents a greater threat, and this results in more vigilance and exploratory behaviour to locate a potentially dangerous intruder. Consistent with this interpretation, both red deer and koalas are more attentive and invest more effort into vocal responses when presented with calls that have lower formants indicative of larger rivals (Charlton et al., 2013; Reby et al., 2005), and howler monkey roaring appears to be important for assessing and repelling opponents in the context of resource defence (Bolt et al., 2019; Sekulic, 1982). It is also possible that females have mating preferences for larger males, and use formants as size cues to preferentially select these individuals as mating partners (Charlton, Ellis, et al., 2012; Charlton et al., 2007). Future studies should examine how size-related formant information propagates in the black howler monkey’s natural environment to determine the distances over which males could still be reliably classified as small, medium or large by receivers (Charlton et al., 2018; Charlton, Reby, et al., 2012). Additional playback studies could also explore whether black howler monkeys alter the acoustic structure of roars in response to different size variants, and test whether oestrous females preferentially approach playbacks of male roars simulating larger callers. It would also be interesting to use play back experiments to examine the degree to which non-linearities, such as deterministic chaos, represent important acoustic components of the highly chaotic howler monkey roars (Muir et al., 2025).

In conclusion, our results indicate that the formant spacing in black howler monkey roars is a reliable cue to body size that is used to assess the threat potential of male conspecifics. They are also consistent with other work that has shown how ‘exaggerated’ vocal traits can still convey reliable information within a given species-specific communication system (Charlton et al., 2011; Harris et al., 2006; Reby & McComb, 2003). Recent comparative work indicates that formants play an important role as cues to body size and identity across mammal vocal communication systems (Charlton & Reby, 2016), and the very low *f*_o_ and/or broadband noise of male black howler roars makes these calls particularly well suited for highlighting formants and increasing the salience of any information encoded by them (Fitch & Hauser, 1995; Ryalls & Lieberman, 1982). Moreover, because mammal vocal signals can potentially act as multicomponent signals of caller characteristics (Candolin, 2003; Koren & Geffen, 2009; van Doorn & Weissing, 2004), multiple additional levels of information, concerning for example, sex, physical condition, social status and hormone levels, may be present in howler monkey roars. Whether male and female conspecifics perceive and attend to this additional information, and how it interacts with body size information, are key questions for future research.

## Acknowledgments

We are grateful to the primate keepers and veterinary staff at Twycross Zoo and Port Lympne for their support with the playback experiments. WTF acknowledges support of Austrian Science Fund (FWF) DK Grant “Cognition & Communication 2” (#W1262-B29).

